# Mechanical de-skewing enables high-resolution imaging of thin tissue slices with a mesoSPIM light-sheet microscope

**DOI:** 10.1101/2025.10.29.684106

**Authors:** Steven Moreno, Sharika Mohanan, Ahmed Elnageh, Erin Boland, Lewis Williamson, Camilla Olianti, Leonardo Sacconi, Godfrey Smith, Eline Huethorst, Caroline Müllenbroich

**Affiliations:** School of Physics and Astronomy, University of Glasgow, Advanced Research Centre, 11 Chapel Lane, G11 6EA, Glasgow, UK; School of Cardiovascular and Metabolic Health, University of Glasgow, 126 University Place, G12 8TA Glasgow, UK; Institute of Clinical Physiology, National Research Council (IFC-CNR), Viale Gaetano Pieraccini 6, 50139 Florence, Italy

## Abstract

Optical clearing combined with light-sheet microscopy enables high-resolution imaging of extended tissue at scale. However, standard mesoSPIM systems are optimised for intact organs and are not suited to thin tissue slices. We present a no-shear imaging protocol using obliquely mounted samples held between refractive-index-matched slides in a 3D-printed frame. This enables mechanical de-skewing during acquisition, minimising post-processing requirements. We demonstrate feasibility in fluorescent bead phantoms and rabbit heart tissue, achieving a 4.8× reduction in processing time and a 1.5× improvement in axial resolution compared to the shear-method. The no-shear method extends mesoSPIM’s utility to fragile, laterally extended tissue sections.

## 1. Introduction

Understanding the correlation between the structure and function of biological tissues requires high-resolution imaging across large spatial scales. Mesoscale light-sheet fluorescence microscopy (LSFM), particularly mesoSPIM-type systems [1, 2], has emerged as a powerful modality to image intact organs and large tissue sections with cellular resolution. Combined with tissue clearing methods [3–5], LSFM has transformed research in developmental biology, neuroscience, pathology and cardiovascular research by enabling comprehensive three-dimensional (3D) reconstructions of complex tissues [6–13].

In large animal models, however, the feasibility of whole-organ clearing and imaging is fundamentally limited by processing time and mounting constraints. In cardiac biology, where model systems aim to replicate human myocardial architecture and pathophysiology, rodent hearts often lack sufficient physiological relevance. However, the rabbit has a comparable electrophysiology to that of humans [14] and is therefore increasingly employed. Yet, due to the prolonged duration and limited penetration efficiency of clearing protocols like Clarity, it is often necessary to isolate and image discrete regions of interest rather than the entire heart. One of the key biological applications that would benefit from mesoscale 3D imaging techniques is the investigation of cardiac remodelling following myocardial infarction (MI). Structural changes in myocardial architecture—including fibrosis, hypertrophy, and alterations in sympathetic innervation—occur over millimetre-scale regions and are not readily captured by conventional 2D histological methods. To model these processes in a translational context, the rabbit heart offers a valuable intermediate between rodent and human physiology. In particular, the percutaneous coronary artery occlusion model in rabbits closely recapitulates aspects of human MI and enables controlled induction of infarction followed by reproducible structural remodelling without the formation of epicardial adhesions [15]. This makes it an ideal system for meso-scale imaging studies aimed at resolving regional remodelling dynamics with cellular resolution. Moreover, many cardiovascular studies are conducted on acute cardiac tissue slices, particularly in large animal and human models, where intact organ imaging is impractical; being able to subsequently clear and image these same slices with light-sheet microscopy would provide valuable structural context to complement prior functional or electrophysiological studies, underscoring the demand for the mesoSPIM’s adaptation for mechanically unsupported, thin tissue sections.

While standard mesoSPIM imaging setups are optimised for whole-organ imaging, where specimens are embedded in a refractive index-matched medium inside a cuvette, adapting these techniques to thin cardiac slices, such as vibratome-cut sections, presents unique challenges. In contrast, inverted Single Plane Illumination Microscopy [16] and Oblique Plane Microscopy [17, 18] configurations allow for direct, horizontal placement of tissue sections on a microscope stage, simplifying their imaging, albeit at the cost of tilted detection geometries that intrinsically produce skewed data sets. Imaging with such systems necessitates computational de-skewing methods [19, 20] to digitally correct for spatial distortions. These approaches can introduce interpolation artifacts [21], increased processing time and higher data storage requirements. Mechanical de-skewing approaches have been described [22], however, these methods have not yet been expanded to mesoSPIM-type light-sheet microscopes. To expand mesoSPIM’s capabilities for imaging laterally extended tissue slices, an alternative approach is required.

In this study, we introduce a custom mounting and optical de-skewing method that enables high-resolution imaging of laterally-extended cardiac tissue slices using a mesoSPIM LSFM. Cleared tissue sections are mounted between refractive index-matched quartz slides held in a 3D-printed frame, which is positioned at a 45° angle between the illumination and detection axes. During acquisition, the sample is translated obliquely through the light sheet via sequential lateral and axial stage movements, resulting in a shear-free image stack with the specimen consistently centred within the field of view. This no-shear acquisition strategy minimises data size and the need for computational de-skewing, significantly reducing processing time while improving axial resolution. We note that this method does not eliminate computational shearing entirely; it performs mechanical compensation during acquisition to pre-align the sample with its coordinate system, thereby greatly simplifying subsequent processing.

We validate this method in three key experiments. First, we compare the spatial resolution of the system across different mounting strategies. We then quantifying improvements in data processing speed and storage requirements for no-shear versus shear acquisitions. Finally, we validate the no-shear approach by imaging a 0.5 mm-thick ventricular section from a rabbit heart, stained for nuclei and cell membranes. This approach provides a biologically translatable platform for exploring the structural basis of post-infarction structural remodelling, a process critical to understanding arrhythmogenesis and functional recovery. Importantly, retaining the three-dimensional integrity of the sample permits the extraction of volumetric data that would otherwise require hundreds of serial histological sections.

Our findings demonstrate that the no-shear method improves axial resolution, reduces computational overhead, and enables high-quality volumetric imaging of structurally delicate tissue sections. This approach extends the mesoSPIM’s utility beyond whole-organ imaging, providing a practical solution for studying thin, laterally extended biological specimens with minimal post-processing artifacts in cardiovascular and other biomedical research contexts.

## 2. Materials and methods

### 2.1. Sample preparation

All animal experiments were approved by the British Council for Animal Research and were conducted in accordance with the UK Animals (Scientific Procedures) Act 1986 and guidelines from Directive 2010/63/EU under Project Licence (PP5254544). All animals were kept and treated in compliance with the local regulations for animal welfare.

#### 2.1.1. Rabbit model of MI

New Zealand White rabbits (3-4 kg) were divided in MI and Sham groups. MI was induced by permanently occluding the main left coronary artery through the percutaneous route [15]. In brief, vascular access was obtained by first dissecting the right carotid artery free, followed by insertion of a 4F sheath into the carotid artery. A 1.5 F microcatheter and 2 mm radiopaque tip were railroaded over a 0.008” guide wire and directed into the left main coronary artery using a 4 F catheter under fluoroscopic imaging. Once the tip was placed in the desired location within the coronary artery, the guide wire, microcatheter and 4 F catheter were retracted. Sham animals underwent the same procedure except for placement of the tip, preventing permanent occlusion [15]. After 6-8 weeks, hearts were excised and used for Langendorff experiment.

#### 2.1.2. Slicing

Following Langendorff perfusion protocol completion, the left anterior free wall of the myocardium was dissected from the rabbit hearts. The tissue collected from the post-MI group was taken to include healthy remote, border zone, and scar regions to enable visualisation of the transition between infarct and healthy myocardium. The tissue was submersion fixed in 4% paraformaldehyde (PFA) and then mounted in a 10% agarose block to support vibratome slicing (SKU: V-1000R). The tissue block was orientated to obtain transmural slices of myocardium (endocardial to epicardial) which were then transferred to PBS with 0.02% sodium azide for storage.

#### 2.1.3. Clearing

Prior to tissue transformation, cardiac slices were mounted between two microscope slides using a custom-designed 3D-printed holder. The slides were spaced to match the nominal vibratome section thickness (ranging from 0.4 mm to 2.0 mm), ensuring consistent support and structural integrity throughout processing. Tissue clearing was performed using a Clarity-based passive clearing protocol optimised for cardiac tissue [23]. Briefly, samples were incubated at 4 °C for 72 hours with gentle agitation in a hydrogel monomer solution consisting of 4% acrylamide, 0.05% bis-acrylamide, and 0.25% VA-044 initiator in 0.01 M phosphate-buffered saline (PBS). The hydrogel solution was injected directly into the sealed tissue mounts to fill the internal chamber, ensuring complete immersion and eliminating trapped air bubbles. Each mount was then placed into a sealable glass jar filled with additional hydrogel solution, which was tightly closed and stored at 4 °C for continued incubation. After 72 hours, jars were slightly opened and transferred to a vacuum degassing chamber (Jeiotech F42400-2121). Atmospheric oxygen was replaced with nitrogen gas to facilitate polymerisation, after which the jars were resealed and incubated at 37 °C for 3 hours to initiate hydrogel cross-linking. Following polymerisation, samples were removed from the jars, slide mounts and immersed in 30 mL of clearing solution comprising 200 mM boric acid and 4% sodium dodecyl-sulfate (SDS) titrated to pH 8.5. Samples were incubated at 37 °C with continuous shaking prevent precipitation of SDS crystals within the tissues for approximately six months, or until optical transparency was achieved. Clearing solution was refreshed three times weekly to maintain detergent efficacy throughout the clearing period.

#### 2.1.4. Staining

Immunolabelling protocols for anti Tyrosine Hydroxylase (anti-TH) and wheat germ agglutinin (WGA) have previously been described in optically cleared pig myocardium and has been applied in this study to prepared rabbit post-MI myocardial slices [24]. Cleared slices were incubated with 1:200 anti-TH (Invitrogen MA5-32984) for 3 days and washed with PBS-T 0.1x for 1 day at RT with gentle agitation followed by incubation with 1:100 WGA - Alexa Fluor 488 (Thermo Fisher, W11261) and Goat anti-mouse Alexa Fluor-647 (Invitrogen C20300) in PBS-T 0.1x for 2 days at RT with gentle agitation. The slices were then fixed with 4% PFA in PBS for 15 min and washed in PBS for 5 min three times. Stained and fixed slices were incubated in 5 mL of EasyIndex (Life Canvas Technologies) and allowed to homogenise for 24 hours prior to imaging for complete RI matching.

#### 2.1.5. Sample mounting

Samples were mounted with a small drop of EasyIndex™ Solution (LifeCanvas Technologies, EI-500-1.52) placed on a single polished 76 mm × 26 mm × 1 mm quartz slide (Portmann Instruments, UQ-1081). The cleared tissue was carefully positioned using a paintbrush to ensure it lay flat without trapped air bubbles. A second drop of EasyIndex™ Solution was applied atop the tissue, followed by a second quartz slide. The sandwiched tissue was then carefully inserted into a 3D-printed slide holder. Once fully positioned, fast curing silicone (Picodent®, PIC. 13007100) was injected along the edges of the mount viewing window to form a watertight seal. A custom 3D-printed topper suspended the mount from the *XY Zθ* sample stage using no adhesives or bolts to reduce strain on the quartz glass. A syringe was used to fill the remaining volume with EasyIndex™solution. The topper incorporated a syringe port and an air exhaust port to facilitate simultaneous injection of solution and expulsion of air, preventing pressure build-ups. A lid was then inserted over the injection and exhaust ports to prevent contamination by dust. The lid contained a 5 mm diameter cylindrical top, allowing it to be securely mounted on a standard 0.5-inch post holder (e.g., Thorlabs PH100/M), which was attached to the *XY Zθ* sample stage. The outer cuvette (Portmann Instruments, UQ-1083) had dimensions of 70 × 65 × 70 mm, larger than a typical mesoSPIM cuvette to accommodate the oblique movement of our mount. The front facets of the mount measured 30 mm in width, requiring a minimum diagonal length of 2 × 30 mm within the cuvette to ensure clearance. The outer cuvette was filled with 68% TDE solution (Sigma-Aldrich, 88561) to match the refractive index of the quartz glass and EasyIndex™ Solution (1.46).

#### 2.1.6. Bead samples

For the rectangular sample cuvette, 1 µm diameter Dragon Green fluorescent beads (Bang Laboratories, FSDG004) were suspended in 1% agarose (Sigma-Aldrich, A9414) at a 1:4000 dilution. The agarose solution was prepared by dissolving low-melting agarose in distilled water, heating it to 70 °C, and allowing it to cool to 40 °C before mixing with the bead suspension. The mixture was then poured into the inner cuvette, solidified at RT, and submerged in 68% TDE (2,2’-thiodiethanol, Sigma-Aldrich, 88561) solution overnight to achieve refractive index matching [4]. For the sandwich mount, the quartz slides (Portmann Instruments, UQ-1081, 76 mm × 26 mm × 1 mm) were inserted into a 3D-printed slide holder (design files available on GitHub [25]) printed with Polylactic Acid (Verbatim # 55317 Natural 1.75mm PLA Filament printed on a Original Prusa MRK 4 3D printer with PrusaSlicer ver 2.9.2 software). Curing Silicone (Picodent®, PIC. 13007100) was applied along the edges to create a watertight seal and was left to set for 24 hours. The Dragon Green bead solution was mixed in 1% agarose at a 1:4000 dilution and injected into the mount using a 0.4 µm diameter syringe, needle gauge 27G. The mount was filled halfway, and a mount topper was placed to seal the chamber. The solution was left to cool and gelatinise at RT for 15 minutes. Once set, the remaining space was filled with 68% TDE solution, and the sample was left to refractive index match overnight before imaging. PSFJ [26] was used to fit PSFs to the bead images and extract lateral and axial Full width half maxima for each of the shear and no-shear datasets.

### 2.2. Imaging

#### 2.2.1. Mesoscale light-sheet microscope

The custom-built light-sheet microscope follows the benchtop mesoSPIM design [2]. Briefly, a laser engine (Omicron LightHub Ultra) enables selection of excitation wavelengths (Luxx CW diode laser, 405 nm, 120 mW; LuxX CW diode laser, 488 nm, 100 mW; OBIS diode-pumped solid state laser, 532 nm, 150 mW; LuxX CW diode laser, 647 nm, 140 mW) with fibre-coupled output. The laser beam undergoes expansion before passing through an electrotuneable lens (ETL, Optotune, EL-16-40-TC-VIS-5D-1-C) for axial scanning of the light sheet [27]. The ETL aperture is imaged onto a galvo scanner (Thorlabs, GVS211), which generates the light sheet. The scanner is positioned at the back focal plane of a modified Nikon camera lens (AF-S 50 mm f/1.4 G), acting as an excitation objective with an f-theta lens configuration. The galvo scanner has a beam diameter of 10 mm, corresponding to an excitation numerical aperture (NA) of 0.1. Fluorescence emission is collected by a 5× and 2× objective (Mitutoyo, 59-876, *f* = 40 mm, NA=0.14 and 59-875, *f* = 100 mm, NA=0.055) and imaged by a tube lens (*f* = 200 mm, Mitutoyo, 54-774) onto the sensor of a large-format scientific complementary metal–oxide–semiconductor (sCMOS) camera (Teledyne Photometrics, 01-Kinetix-M-C). Fluorescence is selectively detected with bandpass and longpass filters (Semrock BrightLine FF01-445/45-25, FF01-514/30-25, FF01-575/59-25 and Chroma ET655LP). The camera has a 29 mm diagonal sensor size, providing a field of view of 5.8 mm. The ETL-driven axial scan is synchronized with the rolling shutter mode of the sCMOS camera. The sample is suspended from an *XY Zθ* stage (Physik Instrumente, 2x L-509.20DG10, 1x L-509.40DG10, ± 100 nm unidirectional repeatability 1x M-060.DG, ±25 µrad unidirectional repeatability) for volumetric image acquisition and the detection focus is actuated by a precision translation stage (Physik Instrumente, M-406.4PD, ±100 nm unidirectional repeatability).

#### 2.2.2. Image acquisition

The image stack of beads in the rectangular cuvette is obtained by translating the sample in steps of 1 µm in the *z* direction. In the sheared method, the sample is translated in *z* with a step size of 1 µm. In the no-shear method, the sample is stepped in *x* and *z* with a step size of 1 µm for an oblique scan in the *x*′ direction. The tissue slices were imaged in the oblique scanning geometry implementing mechanical de-shearing with a *z*-step size of 3 µm. The image stack was then processed through cropping and rotation, followed by an affine transform to map pixels into the correct coordinate system. The images from the shearing scan only require rotation and scaling transformations, while the images from from the no-shear method require shearing and scaling transformations:

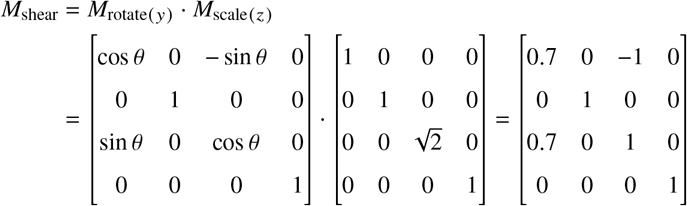

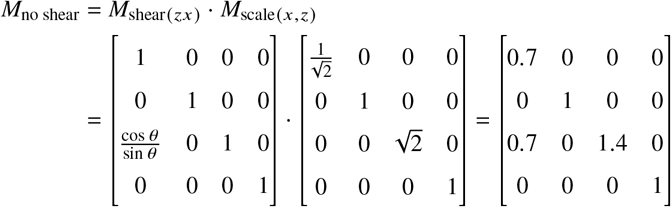

## 3. Results

### 3.1. No-shear approach for mechanical de-skewing of data sets

In conventional LSFM imaging, a cleared tissue section or a whole organ is placed within an inner cuvette, which is immersed in a refractive index-matching liquid inside a larger outer cuvette (Fig.1a). Typically, mounted samples measure approximately 1× 1 × 1 *cm*^3^. The uniform refractive index across both cuvettes ensures minimal optical aberrations as the inner cuvette is scanned through the light sheet for volumetric imaging. However, laterally extended tissue slices, for example thin vibratome-cut section, lack structural rigidity and are difficult to mount without mechanical support. To address this, we developed a custom mounting approach (Fig.1b-d), where the sample is sandwiched between quartz slides held in a 3D printed frame (Fig.1c,d). The slide holder maintains the correct spacing between the quartz slides that corresponds to the nominal thickness of the vibratome cut. The mount is rotated 45° relative to the detection axis, resulting in an oblique scanning geometry (Fig.1a,inset).

**Fig.1.**
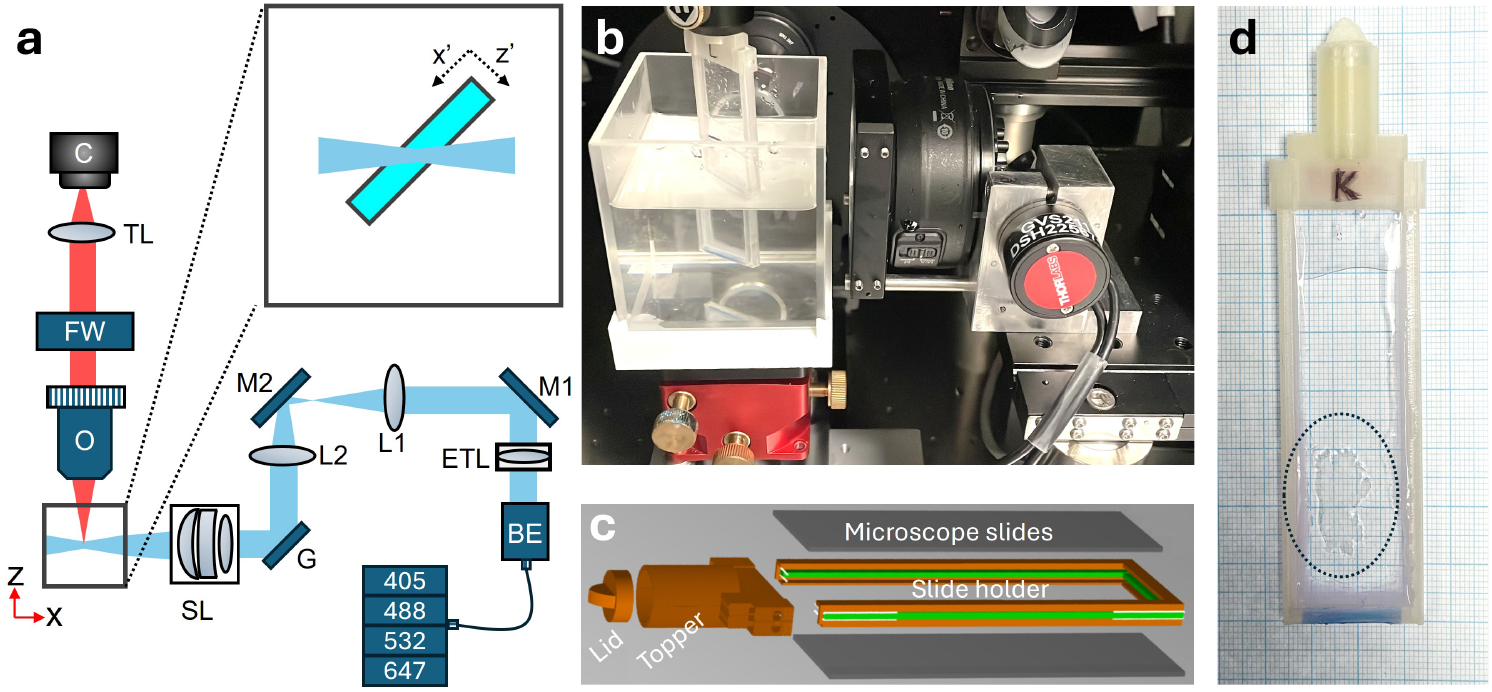
Orientation and mounting of thin tissue slices in the mesoSPIM geometry. **a)** Schematic of the mesoSPIM LSFM. BE: beam expander; ETL: electrically tunable lens, M1,M2: mirrors; L1,L2: lenses; G: galvanometric mirror; SL: scan lens; O: objective; FW: filter wheel; TL: tube lens, C: camera. Inset: orientation of the thin tissue section within the sample chamber. Laser engine provides 405 nm, 488 nm, 532 nm and 647 nm excitation wavelengths. **b)** Photograph of the mount orientation within the outer cuvette. **c)** Exploded view of computer-assisted design of the custom-made slide holder. Spacing between slides (green) corresponds to nominal thickness from vibratome cut. **d)** Photograph of mounted transmural tissue slice; epicardium on the left-hand side and endocardium on the right-hand side.

In LSFMs with oblique scanning geometries, such as iSPIM or OPM, volume reconstruction typically requires computational shearing to correct for the sample’s orientation in the laboratory coordinate system (here denoted with *x, y*) [21, 28]. In these methods, *z*-scanning (Fig.2a) inherently produces a sheared volume that must be computationally de-sheared. This process necessitates pixel padding of the data volume (grey areas) increasing its file size. Finally, an affine transformation, requiring additional processing time and padding, maps pixels into the correct sample coordinate system (*x*′, *z*′). In sheared volume reconstruction, this affine transformation involves scaling and rotation (See Supplement for supporting content). Also for our laterally extended samples mounted at 45°, *z*-scanning inherently produces a sheared volume resulting in increased data size and processing time. To address this, we introduce an optimised volume reconstruction methodology, called “no-shear” method, by obliquely scanning the sample through the light sheet (Fig.1b).

**Fig.2.**
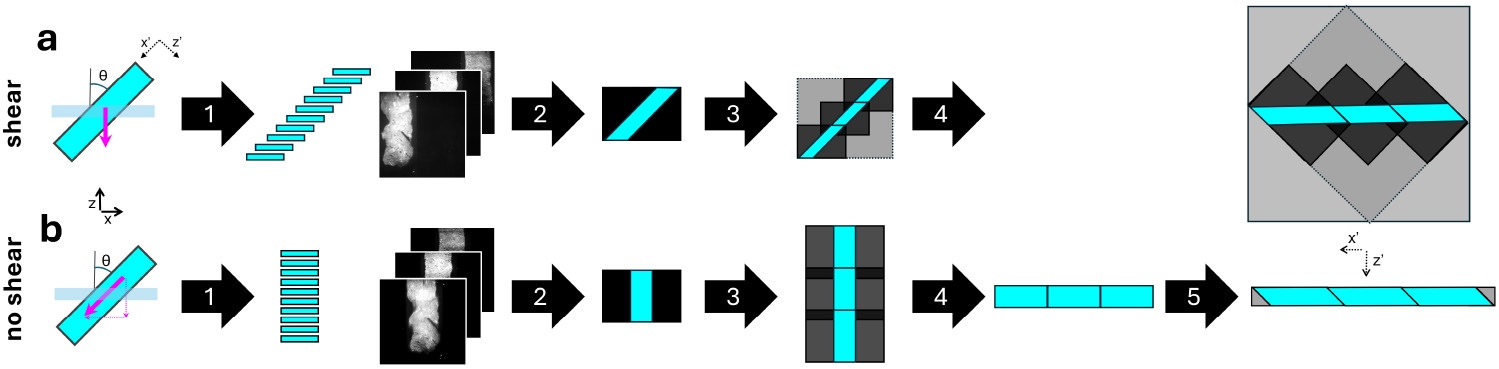
Oblique scanning geometry and volume reconstruction for the **a)** shear and **b** no-shear method. Stage scanning direction (magenta arrows) are indicated with respect to the laboratory coordinate system (*x, z*). The volume reconstruction produces a stack orientated in the sample coordinate system (*x*′, *z*′). Shear method: 1) Sample acquisition produces sheared raw data stack in which the sample translates from left to right with each image. 2) Padding with empty pixels (grey area). 3) Tiled acquisition requires further padding to stitch sample volume along *x* and *z*-direction. 4) Computational shearing involves rotation and additional padding. Cropping of dataset is only possible after this process. No-shear method: 1) Sample acquisition produces raw data stack in which the sample remains in the central part of the field of view (2). 3) Stitching is only required along *z*-direction. Unlike the shear method, no further padding is needed. 4) Rotation of the data stack by 90 degree. Pixels without content can be cropped to reduce data size. 5) Scaling along *x*′ and shearing along *z*′ of the data stack reconstructs the sample volume.

We compensate for shear mechanically by introducing a depth-dependent lateral shift using the *x*-stage. The shift required for each slice in the *z*-stack is given by: *x*_*s*_ = *nz*_*s*_ tan *θ* where *θ* is the angle between the sample and the detection axis, *x*_*s*_ is the shift in the x -direction to re-centre the image on the camera for the *n*th slice and *z*_*s*_ is the z step size. For our system *θ* = 45° and the equation simplifies to *x*_*s*_ = *z*_*s*_ allowing straightforward implementation. In the no-shear data sets acquired in this way, the tissue slice always remains in the central part of the field of view.

Once the no-shear dataset is acquired, a few simple steps with minimal computational overhead are applied to map pixels accurately into the correct sample coordinate system. Firstly, since the tissue occupies only the central subset of the camera frame, images are easily cropped to minimise data size. Secondly, we perform a 90° rotation. This operation, reorients the dataset along the *x*′ plane, enabling immediate visualization. Finally, scaling along *x*′ and shearing along *z*′ reconstructs the sample volume.

### 3.2. System characterisation

First, we quantified the spatial resolution of our LSFM employing the standard rectangular cuvette to benchmark the performance of mesoSPIM. The violin plots (Fig.3a) demonstrate a spatial resolution that is consistent with published literature for the mesoSPIM light-sheet microscope [1, 2]. The mean full-width at half-maxima (FWHM) of sub-resolution fluorescent beads in the *xyz* directions for the standard cuvette were (4.01 ± 0.85) µm, (5.27 ± 1.05) µm, and (5.56 ± 0.72) µm, respectively (number of beads, n = 196). Note that these sizes are reported in the laboratory coordinate system (*x, y, z*).

**Fig.3.**
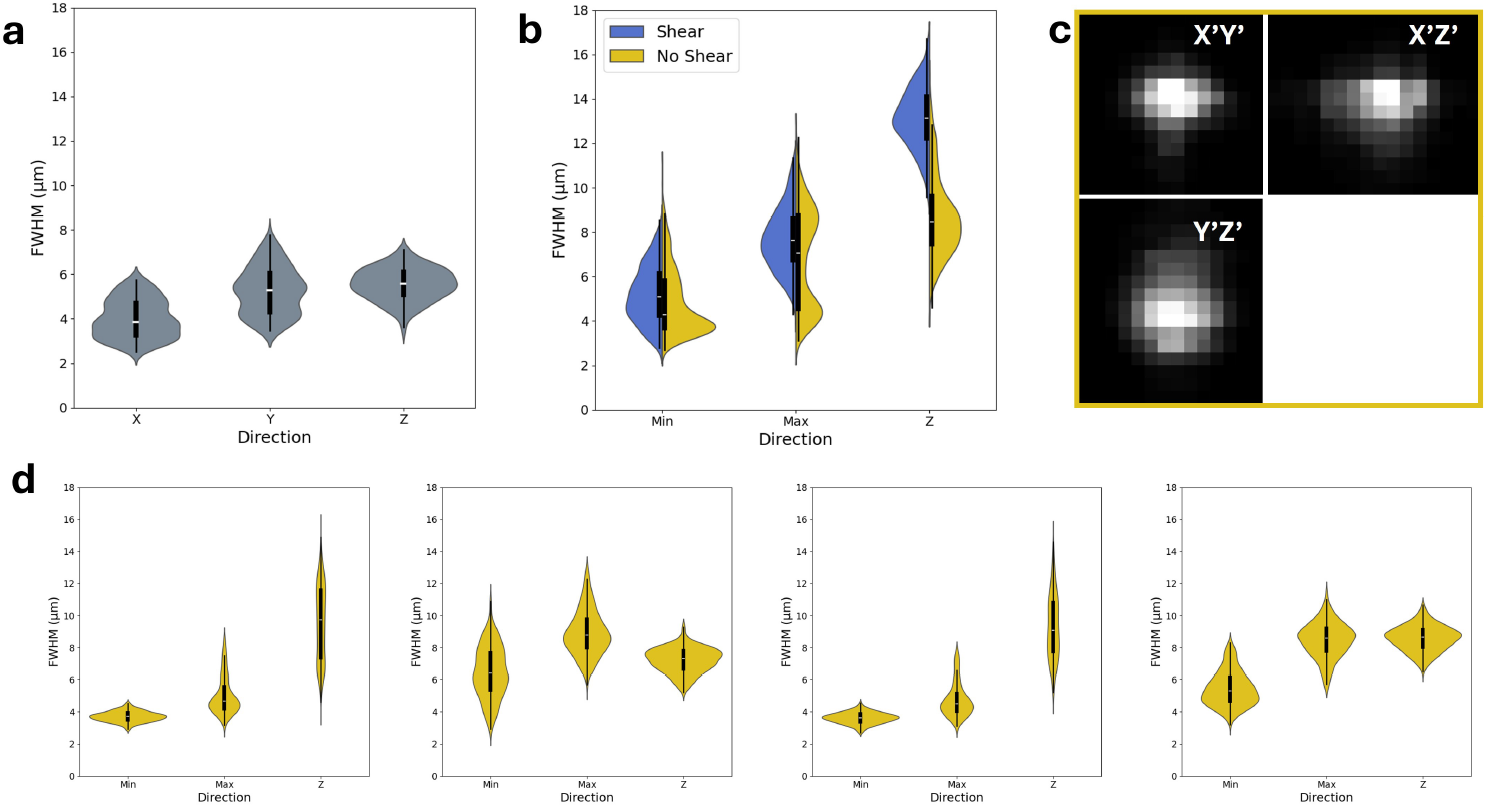
Point spread function analysis of 1 µm diameter fluorescent beads. a) Violin plots of the full width at half maxima (FWHM) of beads (*n* = 196) embedded in agar within the rectangular cuvette. Note that these sizes are reported in the laboratory coordinate system (*x, y, z*). b) Violin plots comparing the FWHM of beads embedded in agar in sandwich mounts using the shear (blue, *n* = 562 beads in 1 mount) and no-shear method (yellow, *n* = 1236 beads in 4 mounts). Note, dimensions are given in the sample coordinate system (*x*′, *y, z*′). An *R*^2^ threshold of 0.95 ensured quality of fit. White line marks the median, black box marks the 25th and 75th percentiles. Whiskers mark out 1.5 times the inter-quartile range. c) Qualitative view of point spread function (no-shear) visualises astigmatism present. d) Violin plots for four different 1 mm sandwich mounts quantify variability introduced by the sample holder.

We then compared the resolution achieved when mounting the sample at 45° using the shear and no-shear methods (Fig.3b). When comparing these values to those obtained in the standard rectangular cuvette, it is important to note that the values reported for the shear and no-shear methods are given in the sample coordinate system (*x*′, *y, z*′). The violin plots for the shear method (blue, left) exhibit worse resolution in all dimensions but particularly in *z* compared to the cuvette. This is likely due to aberrations caused by the oblique mounting and the custom-made nature of the sample holder. In contrast, the no-shear method violin plots (yellow, right) showcase an improved resolution in all dimensions with respect to the shear method. For the shear method, we obtained a mean spatial resolution in the *xyz* directions of (5.22 ± 1.27) µm, (7.70 ± 1.39) µm, and (13.15 ± 1.36) µm, respectively (n = 562 beads). For the no-shear method, these values were (4.88 ± 1.51) µm, (6.82 ± 2.25) µm, and (8.72 ± 1.80) µm, respectively (n = 1236 beads). As expected, we observed astigmatism due to non-uniform stress on the quartz slides, leading to asymmetry in the lateral dimensions (Fig.3c). The anisotropy ratio, 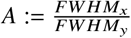, is 0.72 for both modalities indicating that there is no change in the ratio of the lateral minimum to maximum bead size. We further observe a significant axial resolution enhancement in the no-shear method, which is evident from the lower median albeit at increased spread in the axial FWHM distribution. We attribute the 1.5× reduction (13.15 ± 1.36) µm reduced to (8.72 ± 1.80) µm) in axial resolution compared to the shear method to the ability to fine-tune ETL parameters. The ETL sweep is confined to the sample region at the centre of the camera’s field of view, as the tissue slice remains stationary in the centre enhancing axial resolution. To assess the reproducibility of our custom 3D-printed mounts, we quantified the point-spread function across four 1 mm sandwich mounts (Fig. 3d). The overall resolution remained consistent across mounts, indicating robust optical alignment and mounting reproducibility. While small mount-to-mount variations were observed, primarily in the lateral (*x, y*) dimensions, the axial (*z*) resolution showed minimal deviation, confirming that the sample holder design preserves the optical quality along the detection axis. This suggests that small mechanical tolerances in the 3D printing or stress in the quartz slides may lead to mild lateral astigmatism, but these effects do not significantly compromise axial resolution or imaging fidelity.

#### 3.3. Computational overhead of volume reconstruction

To assess the computational efficiency of the shear and no-shear methods, we analysed the final size of the stitched data set and processing time needed for volume reconstruction. A 2×2 tiled dataset was acquired from a cleared left ventricle slice of rabbit heart with the the shear and no-shear method.

In the sheared method, tiling requires a 10% overlap in both the *x* and *z* dimensions requiring customisation of the tiling process from conventional methods that tile only in *x* and *y* (see Fig.2). This redundancy increases the raw dataset size and necessitates additional processing steps to merge overlapping regions. Furthermore, the datasets obtained with the shear method cannot be cropped, as the skewed acquisition results in non-trivial empty spaces within the imaging volume. Consequently, the final stitched file is substantially larger than the raw dataset (Fig.4a), increasing storage demands. The normalised increase in stitched file size for the no-shear method is by a factor of 4.24 ± 0.07 (Fig.4b). Conversely, the no-shear method eliminates the need for tiling in the *x*′ dimension, as the scan is performed along the oblique axis with the sample size constrained only by the working distance of the detection objective. This minimises redundancy in overlap and allows for cropping of empty pixels, further minimising the dataset size (Fig.4a). In the no-shear method, the size of the stitched data set is actually smaller than the raw data by a factor of 0.75 ± 0.004 (Fig.4b). This constitutes a reduction in stitched data size of factor 5.63 ± 0.10 of the shear method compared to the no-shear method.

**Fig.4.**
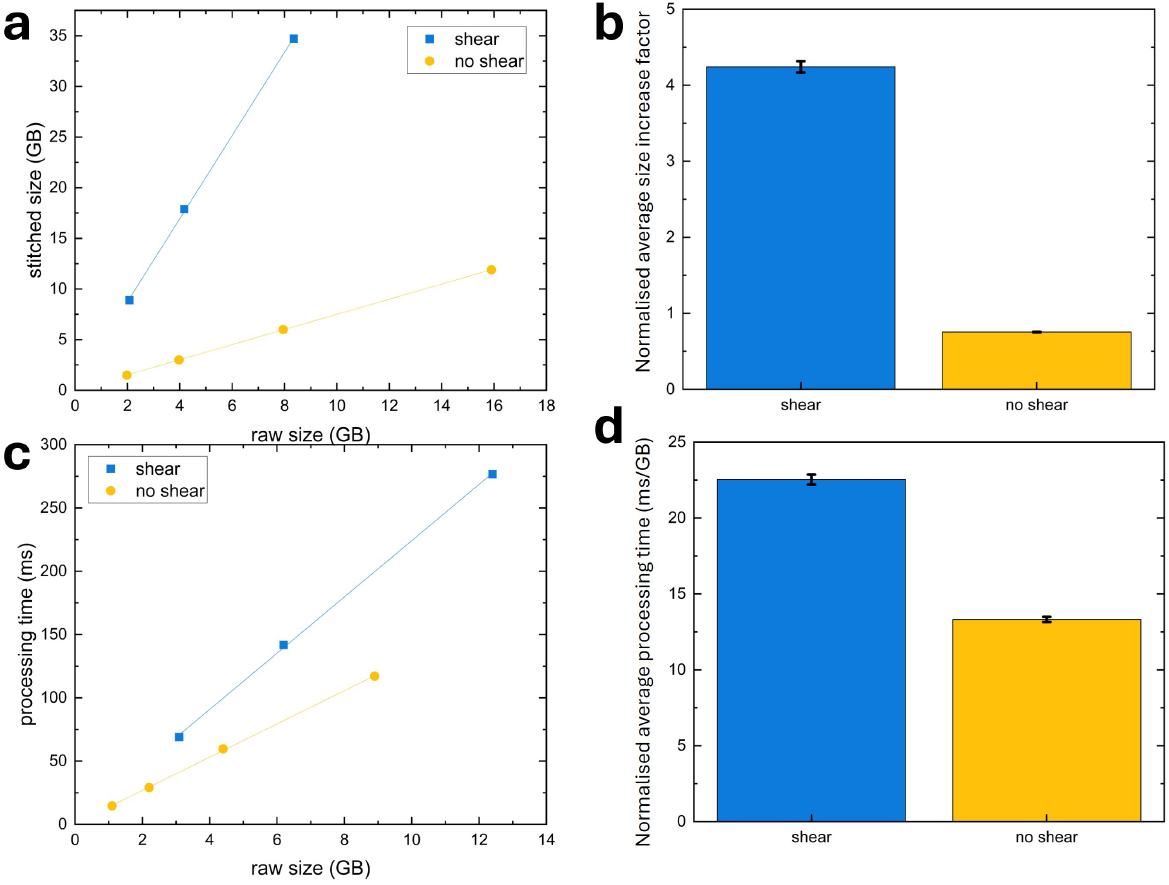
Computational overhead of volume reconstruction. **a)** Final data size after de-skewing as a function of raw file size for the shear method (blue squares) and the no-shear method (yellow circles). **b)** Average data file increase. Error is standard deviation. **c)** Processing time for the affine transformation as a function of input file size for the shear method (blue squares) and the no-shear method (yellow circles). **d)** Average processing time (ms/GB). Error is standard deviation.

Due to the smaller file size, computational processing time required for the affine transformation is consistently lower when using the no-shear approach (Fig.4c). The normalised processing time for the shear method is (22.53 ± 0.33) ms GB^−1^ (Fig.4d). Conversely, the no-shear method requires a normalised processing time of (13.32 ± 0.18) ms GB^−1^. This consistent reduction by a factor of 1.69 ± 0.03 in processing time outperforms the shear approach, highlighting the suitability of the no-shear method for high-throughput imaging of tissue slices, where large volumes must be processed efficiently without excessive computational burdens.

### 3.4. Laterally extended tissue imaging

We finally demonstrate the utility of the oblique mounting and no-shear scan method by imaging a laterally-extended tissue slice. Using the no-shear method, we imaged a 2 mm-thick Clarity-cleared tissue section from the left ventricle of a healthy rabbit heart (Fig.5a) stained with Sytox Green to label nuclei. Resolution is sufficient to identify individual nuclei across the tissue slice as highlighted by three regions of interests (ROIs) (Fig.5b-d). Next, we imaged a 0.5 mm-thick Clarity-cleared tissue section from the left ventricle of a healthy rabbit heart (Fig.5e). To visualize relevant structures, the tissue was double-stained with WGA conjugated to Alexa Fluor-488 to label cell membranes and DAPI to label nuclei (Fig.5a). The no-shear method yields sufficient contrast and resolution to identify individual nuclei and delineate membrane boundaries throughout a 500 µm-thick myocardial section highlighting uniform visibility of subcellular structures across the tissue depth in dense, fibrous myocardium. We identified nuclei in the DAPI channel and extracted their sizes in the sample coordinate system (*x*′, *y, z*′) across two representative ROIs (Fig.5f,g). Measurements of nuclear size in the no-shear datasets revealed dimensions consistent with the expected range for cardiomyocyte and fibroblast nuclei, typically spanning 5 µm to 15 µm. The nuclei were generally elliptical rather than perfectly spherical, as expected due to their intrinsic morphology and anisotropic organisation within myocardial tissue. These 3D data suggest that robust nuclei segmentation is possible due to high axial resolution and minimal distortions of the no-shear technique. This finding opens up the possibility to quantitatively study regional heterogeneity in nuclear morphology as a biomarker in cardiac pathology. The axial FWHM of identified nuclei is plotted across the two regions of interests (Fig.5h) alongside the axial resolution limits of both shear and no-shear methods. Many nuclei would not have been resolved with sufficient contrast using the shear method as FWHM of nuclei approaches or falls below the resolution limit. The enhanced axial resolution of the no-shear method enables volumetric segmentation of individual nuclei that would otherwise appear blurred or merged. The resolution improvement achievable with the oblique mounting and no-shear acquisition method therefore allows for the segmentation of a larger percentage of nuclei than would have been the case with the shear method. The nuclear size distributions observed across the two regions of interest illustrate the method’s ability to resolve and quantify subcellular morphology throughout a 500 µm-thick myocardial slice. The improved axial resolution of the no-shear method ensures that even densely packed or elongated nuclei are faithfully segmented, enabling comparison of nuclear morphology between myocardial regions. This capability is particularly valuable in the study of post-MI remodelling, where nuclear hypertrophy or elongation may correlate with fibrosis, cardiomyocyte stress, or immune cell infiltration.

**Fig.5.**
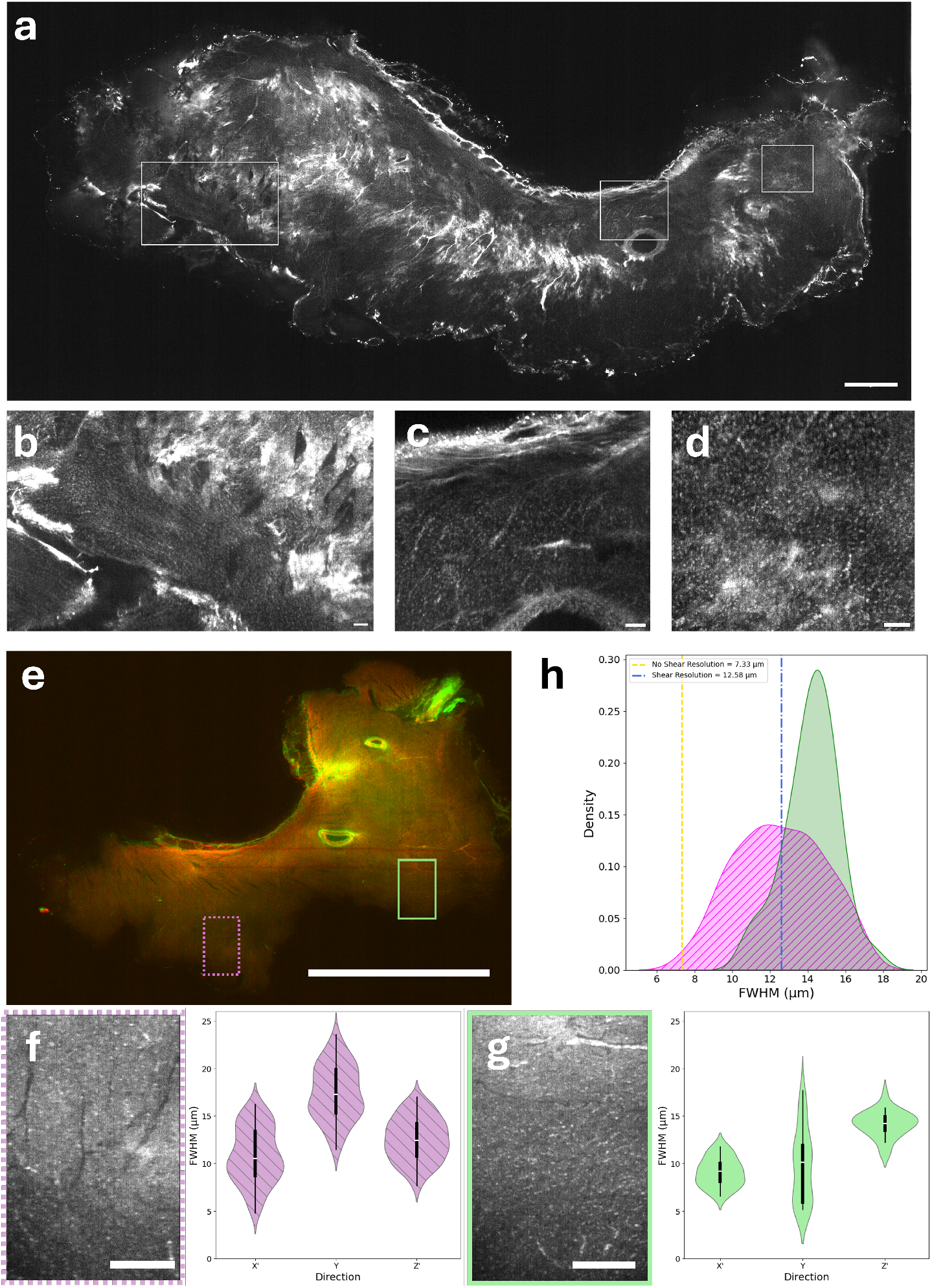
**a)** 2 mm-thick ventricular section of rabbit heart cleared with the Clarity protocol and stained with Sytox Green. Scalebar 1 mm. **b-d)** Regions of interests, scalebar 100 µm. **e)** 500 µm-thick ventricular section of rabbit heart cleared with the Clarity protocol and stained with WGA-AF488 and DAPI. **f**,**g)** Resolution was sufficient to analyse nuclei size across regions of interest (green and magenta ROIs, scalebar 300 µm). Nuclei size distribution is reported for both ROIs in the sample coordinate system (*x*′, *y, z*′). **h)** FWHM of the axial size of segmented nuclei in the two ROIs. Also indicated are the axial resolutions of the shear and no-shear methods as determined from the bead analysis. The enhanced axial resolution of the no-shear method compared to the shear-method is essential to distinguish nuclei across the magenta ROI.

As a further example, we imaged a 500 µm-thick tissue section from the left ventricle of a rabbit heart (Fig.6a) cleared with the Clarity protocol. To visualize relevant structures, the tissue was double-stained with WGA-AlexaFluor488 ((Fig.6b)) to label cell membranes and immunolabelling for anti-Tyrosine Hydroxylase (anti-TH, Fig.5c) to highlight sympathetic neuronal structures. The preservation of structure throughout the 500 µm-thick myocardial slice suggests effective penetration of clearing agents, fluorescent stains and light, supported by the improved optical quality of no-shear imaging. Neuronal remodelling post-MI is well-documented but technically challenging to study in 3D. While light-sheet imaging has been used to study cardiac innervation patterns in whole mice hearts [29], no prior study has leveraged the mesoSPIM to map neuronal distributions in MI-induced heart slices at high resolution in rabbit heart. Our objective was to determine whether the no-shear method enabled 3D reconstructions of spatially continuous delicate features like sympathetic fibres and innervation heads in thick cardiac slices. We found that the innervation network could be reconstructed in selected ROIs (Fig.6d) across the laterally extended tissue slice. The preservation of staining and micro-architecture was confirmed using high-resolution multiphoton microscopy, indicating that tissue integrity was maintained during the clearing and mounting process (Fig.6e). Using oblique mounting and the no-shear method, image resolution was sufficient to distinguish innervation networks and obtain 3D reconstructions of the anti-TH-labelled structures. The antiTH-stained fibres retain morphological continuity and spatial coherence, indicating the no-shear method’s ability to resolve extended, filamentous structures in 3D. Sympathetic innervation in the heart is known to undergo complex remodelling following MI, including regional hyperinnervation, axon sprouting, and nerve degeneration. Although this study does not yet quantify those features, our findings demonstrate that the no-shear method offers sufficient resolution and imaging fidelity to support future quantitative mapping of innervation architecture in post-MI tissue. The qualitative visualisation of sympathetic innervation (Fig.6d) highlights the biological value of the no-shear method for mapping fine neuronal structures in 3D. Tyrosine hydroxylase-positive fibres were consistently resolved across the thickness of the tissue section, with sufficient continuity to support spatial reconstruction. These data suggest that sympathetic axons, which are typically slender and highly branched, can be tracked over extended distances within mesoscopic volumes using our approach. While this study did not quantify innervation metrics, the imaging quality is compatible with future quantitative analysis of innervation density, fibre orientation, and regional remodelling after MI. Crucially, because the optical clearing and mounting pipeline can be applied to tissue that has undergone prior functional experiments, such as electrophysiological studies, it enables retrospective structural mapping of the same samples. This opens the possibility to directly relate functional readouts to three-dimensional structural features within the same tissue, advancing structure–function correlation in cardiac research.

**Fig.6.**
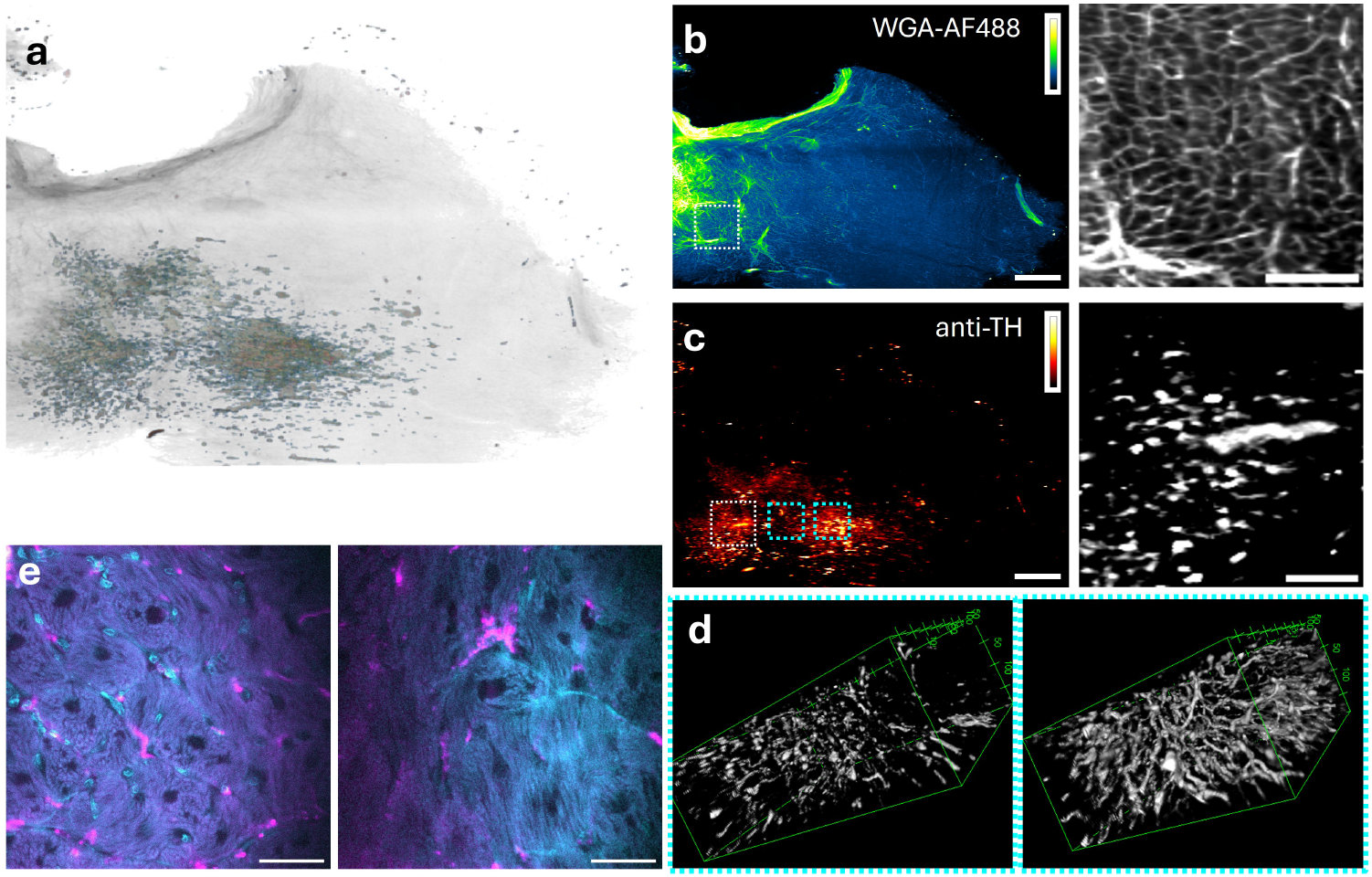
**a)** Overview of a 500 µm-thick ventricular section of rabbit heart cleared with the Clarity protocol and stained with **b)** WGA-AF488 and **c)** anti-TH. Scalebar 1 mm. Right hand side shows zoomed in region of interest (ROIs) indicated. Scalebar 250 µm. Anti-TH signal spatially reconstructed in two ROIs highlighted in cyan in panel (c). High resolution multiphoton microscopy images of the sample micro-architecture. Cyan: WGA-AF488, magenta: anti-TH (scalebar 50 µm).

## 4. Discussion

We developed a no-shear acquisition strategy for mesoSPIM-type microscopes that facilitates volumetric imaging of laterally extended tissue slices. By combining a custom, oblique mounting approach with coordinated scanning in the axial and lateral directions, this method partially compensates for the shear introduced by oblique sample orientation. The resulting raw data remain centered within the field of view and are easily cropped and de-sheared thereby streamlining the post-processing workflow for alignment in sample coordinates. Unlike conventional oblique scanning, which demands computationally intensive affine transformations and introduces interpolation artifacts, our approach simplifies the reconstruction pipeline by reducing the amount of padding and resampling required.

Crucially, our method differs from full mechanical de-shearing techniques reported in other systems. For example, Lin *et al*. [22] implemented mechanical shearing in a custom-built upright diSPIM-like ASLM configuration, enabling data acquisition directly into the sample coordinate system. By contrast, our method, designed for compatibility with existing mesoSPIM hardware [1, 2], does not eliminate computational shearing entirely. Instead, it shifts the sample laterally during axial scanning to partially de-skew the dataset at acquisition. This reduces the volume of empty pixels, streamlines the transformation process, and decreases overall processing time by a factor of 1.69 compared to conventional sheared imaging. Additionally, at 5× magnification, tissue is imaged along its depth in the *z*′ direction with pixel sampling at 1.31 µm in the *x, y* directions. Since the resolution of our microscope ranges from 5 µm to 7 µm laterally, oversampling allows pixel binning in *x, y* without compromising resolution.

Our system differs not only in architecture but also in its practical scope. While Lin *et al*. focus on high-NA, isotropic imaging in an upright configuration, we target a widely used platform optimised for whole-organ imaging. Our method adapts mesoSPIM for mechanically unsupported thin slices, offering a practical solution on pre-existing setups that can be adopted with minimal hardware and software modifications. Our custom mounting system supports reproducible orientation of fragile tissue slices at a 45° angle. This alignment, essential for the no-shear acquisition, enables consistent imaging across experiments. We show that this geometry permits tuning of the ETL over a confined central region of the field of view, which improves axial resolution by a factor of two compared to conventional shear-based acquisition.

One limitation of our approach is the variability introduced by the custom-made oblique tissue mounts. Stress-induced astigmatism observed in some mounts suggests that mechanical tolerances in the quartz slide holders may introduce optical aberrations, particularly in the lateral dimensions. Future iterations of the mounting system could incorporate stress-relief features or alternative materials to mitigate this effect.

Importantly, while we still apply a residual affine transformation to align data into the sample coordinate system (*x*′*yz*′), the computational demands are significantly reduced. Our volume reconstruction requires only rotation and minor rescaling, in contrast to full affine transformations involving scaling, translation, and rotation in multiple axes. Our benchmarking reveals a more than fivefold reduction in stitched file size compared to sheared acquisition and a corresponding improvement in post-processing throughput. These savings become critical when imaging large tissue volumes or executing multi-tile acquisitions. Additionally, mechanically shifting the image reduces interpolation artefacts introduced by computational shearing. Since stitching is not required in the *x*′ direction, stitching artifacts are further minimized. Finally, along the *x*direction, this imaging geometry also minimises shadow and scattering artifacts [9, 30].

We validated the performance of this method using 1 µm fluorescent beads, confirming minimal introduction of detrimental aberrations like astigmatism while demonstrating substantial axial resolution improvements. Application to rabbit cardiac tissue slices demonstrated that fine-scale features such as sympathetic innervation could be reliably detected across millimetre-scale fields of view, underscoring the method’s suitability for structurally delicate tissues. Owing to its compatibility with vibratome-cut slices, this approach is readily transferable to other large-animal and human cardiac preparations including sheep, pig, and human myocardial slices.

## 5. Conclusion

We present a mechanically driven, no-shear imaging strategy for mesoscale light-sheet microscopy that enables high-resolution imaging of fragile, laterally extended tissue slices without requiring major hardware modifications to existing mesoSPIM systems. By eliminating the need for computational de-skewing, this method significantly reduces processing time, minimises interpolation artefacts, and improves axial resolution. Our optical de-skewing approach enhances spatial fidelity and data quality by maintaining the sample within the optimally illuminated region of the field of view during acquisition. It also reduces data volume by eliminating padding pixels, streamlining downstream analysis and storage requirements. This methodology extends the capabilities of mesoSPIM systems beyond whole-organ imaging to include vibratome-cut sections and other mechanically unsupported samples where whole organ imaging is not possible. As an accessible and scalable solution, it offers a practical upgrade path for laboratories seeking to perform high-throughput volumetric imaging with minimal post-processing overhead.

## Acknowledgment

The authors gratefully acknowledge the Cellular Analysis Facility (CAF) for their support & assistance in this work. The authors further thank Dr Nikita Vladimirov for a helpful discussion.

## Contributions

**Steven Moreno**: data curation, investigation, methodology, visualisation, writing -review & editing. **Sharika Mohanan**: conceptualisation, data curation, methodology, software, formal analysis, investigation, visualisation, supervision, writing-original draft preparation. **Ahmed Elnageh**: formal analysis, investigation, software, visualisation, writing -review & editing. **Erin Boland**: investigation, methodology, Validation and writing – review & editing. **Lewis Williamson**: investigation. **Camilla Olianti**: methodology, validation, writing -review & editing. **Leonardo Sacconi**: resources, methodology, validation, writing -review & editing. **Godfrey Smith**: conceptualisation, funding acquisition, methodology, resources, supervision, validation, writing -review & editing. **Eline Huethorst**: conceptualisation, investigation, validation and writing – review & editing. **Caroline Müllenbroich**: conceptualisation, formal analysis, funding acquisition, investigation, methodology, project administration, resources, supervision, validation, visualisation, writing-original draft preparation, writing -review & editing.

